# Molecular mechanism of the key dormancy regulator DosR-dependent transcription activation in *Mycobacterium tuberculosis*

**DOI:** 10.1101/2025.10.11.681782

**Authors:** Jing Shi, Zhenzhen Feng, Han Fu, Yirong Huang, Liqiao Xu, Qian Song, Wei Chen, Yu Feng, Liang-Dong Lyu, Wei Lin

## Abstract

*Mycobacterium tuberculosis* (Mtb) is the etiological agent responsible for the worldwide disease Tuberculosis. DosR is recognized as the key dormancy regulator of Mtb due to its global transcriptional regulation in intracellular adaptation and long-term persistence prevailing in latent infection. However, the molecular mechanism regarding DosR-dependent transcription activation remains inadequately defined. Here, we successfully determined the cryo-EM structure of an intact DosR-dependent transcriptional activation complex (DosR-TAC), comprising Mtb RNA polymerase (RNAP), DosR, and the hypoxic promoter DNA. In DosR-TAC, two DosR monomers symmetrically dimerize through their N-terminal receiver domains (DosR_RECs) and the distinct α10 helices of the C-terminal DNA-binding domains (DosR_DBDs). Unlike its inhibitory configuration, the DosR dimer undergoes significant conformational changes within the original linker, isomerizing DosR_REC into a canonical (βα)5 fold. This extended linker affords the DosR_DBDs and DosR^I^_REC to contact promoter DNA, while concurrently interacting with the conserved domains of RNAP (σ^A^R4, αCTD, and αNTD). Additionally, RT-qPCR and growth curve assays of the wild-type and DosR mutant strains, also unravel the physiological importance of this divergent linker. These findings, together with prior hypotheses regarding DosR, support an ‘allosteric activation-recruitment’ model for DosR. Altogether, these results highlight the molecular mechanism of DosR-dependent transcription activation critical for dormancy survival of Mtb, and offer potential targets for developing anti-dormancy strategies against persistent tuberculosis.

## Introduction

Tuberculosis (TB) remains the leading cause of death attributable to infectious diseases worldwide, continuing to pose a significant global health challenge. According to the World Health Organization, TB accounts for approximately 1.25 million deaths and 10.8 million new infections annually (1). *Mycobacterium tuberculosis* (Mtb), being the primary etiological agent of TB, has evolved diverse transcriptional adaptation mechanisms that enable it to withstand intracellular restrictive environments (2-4). These adaptations facilitate the bacterium’s survival in non-replicating states, thereby contributing to prolonged persistence and the establishment of asymptomatic latent tuberculosis infection (LTBI), characterized by the formation of granulomas (5-7). Epidemiological data indicate that approximately one-quarter of the global population has been infected by Mtb with an occurrence of latent TB in about 90%, among which 5% to 10% may experience reactivation of the infection, particularly when immune function is compromised (8). Despite these insights, the molecular mechanisms governing transcriptional regulation during TB latency remain incompletely understood.

The DosRST two-component system, comprising the histidine kinase DosS (or DosT) and the dormancy survival regulator DosR (also referred to as DevR), is essential to the adaptation and pathogenicity of Mtb during latent infection (3, 7, 9-14). In response to hypoxic conditions, such as NO/CO, DosR is phosphorylated by its cognate kinase, which subsequently increases its DNA binding affinity to specific DosR binding boxes (13, 15-17). Among the most extensively characterized members of these regulons is the alpha-crystallin-like *hspX* gene (18-20). Numerous studies have demonstrated that DosR modulates the expression of approximately 50 genes associated with dormancy development, survival under hypoxia, virulence, and the metabolic reprogramming necessary for the transition of active Mtb into a low-metabolic dormancy state (13, 21). Consequently, DosR is widely recognized as the key regulator of dormancy in Mtb.

Furthermore, the observed sequence similarity and comparable hypoxia response mediated by the DosRS regulatory system in Mtb, *Mycobacterium smegmatis* (*M. smegmatis*), and *M. bovis BCG* imply that the molecular mechanisms underlying the dormancy response are likely conserved across these species (21-24). *M. smegmatis* could be served as a valuable model for investigating the transcription regulation and dormancy survival strategies of their counterpart in Mtb. Intriguingly, the avirulent strain *M. smegmatis* harbors two distinct sets of DosR proteins (MSMEG5244 and MSMEG3944), however, only the first set MSMEG5244 appears to be functionally relevant to hypoxic induction, whereas the second set MSMEG3944 seems to lack functional activity (23, 25). Although sequence and structural alignments reveal similarities among DosR homologues (***SI Appendix*, Fig. S1**), elucidating the critical differences in transcription regulation could provide novel insights into the functional basis governing Mtb virulence during latency induction.

Distinct from other response regulators within the NarL/UhpA subfamily, the self-inhibited Mtb DosR features an atypical N-terminal (βα)4 fold receiver domain (DosR_REC, comprised of residues 1-97 spanning from α1 to α4, β1 to β4), a C-terminal DNA-binding domain (DosR_DBD, comprised of residues 150–217 spanning from α7 to α10), and a long connecting Linker (comprised of residues 98-149 including α5’α6) (***SI Appendix*, Fig. S1**). Additionally, phosphorylation of the DosR_REC at the conserved residue Asp54 or a mutation D54E with similar activation effects is required for the DNA binding activity of DosR_DBD (20, 26, 27). As depicted in the crystal structure, the unphosphorylated full length *Mtb* DosR forms dimers via its linker α5’α6 (20), the DosR_DBD dimers assemble into tetramers through the α7/α8 interface or bind hypoxic promoter DNA in a dimeric configuration, stabilized by the indispensable α10/α10 interface (26, 28). These observations imply that substantial significant structural rearrangements and conformational changes accompany DosR dimerization and DNA binding. Moreover, mutational analyses have demonstrated that the interaction between DosR and the housekeeping sigma factor σ^A^ of Mtb RNA polymerase holoenzyme (RNAP) is also required for the transcription activation of hypoxia-responsive genes, indicative of an interceptable molecular interface for exploring potential inhibitors or interventions (29). However, due to the absence of structural data on the full-length activated DosR and the functional DosR-dependent transcriptional activation complex, the precise mechanism by which Mtb DosR activates and coordinates its DosR_REC and DosR_DBD to cooperatively engage promoter DNA and recruit RNAP remain unresolved and represent critical questions for ongoing investigations.

In this work, we determined the cryo-EM structure of an intact DosR-dependent transcriptional activation complex (DosR-TAC) comprising of Mtb RNA polymerase (RNAP), DosR, and the hypoxic promoter DNA. Within DosR-TAC, two DosR monomers form a centrosymmetric dimer through interactions involving the N-terminal DosR_RECs and the distinct α10 helices from the C-terminal DosR_DBDs. Unlike its inhibitory configuration, the DosR dimer exhibits significant conformational rearrangements in the unique linker helix α5’. This helix undergoes isomerization into the canonical β5 and α5 structural elements, resulting in an extension of the linker helices α6 which spatially separate the canonical (βα)5 folded DosR_RECs from the repositioned DosR_DBDs. This distinctive architecture enables the DosR_DBDs and DosR^I^_REC to interact with promoter DNA while concurrently engaging the conserved R4 domain of σ^A^R4 (σ^A^R4), as well as the C-terminal domain (αCTD) and the N-terminal domain (αNTD) of the α subunit of RNAP, respectively. *In vivo* analyses, including RT-qPCR and growth curve assays of wild-type DosR and its mutants also unravel the functional roles of key residues within the linker region. These findings, together with prior hypotheses regarding DosR, support an ‘allosteric activation-recruitment’ model for DosR. Altogether, these results reveal the conformation flexibility and structural basis of DosR-dependent transcription activation critical for the dormancy survival of Mtb, and present potential targets for developing promising strategies against persistent tuberculosis.

## Results

### Overall cryo-EM Structure of the Mtb DosR-TAC

Considering the *hspX* promoter, a member of the DosR regulon, is well-established as being closely linked to the dormancy and persistence of Mtb (18, 20, 30), it was chosen to explore the structural basis underlying DosR-dependent transcription activation. Initially, we synthesized an *hspX* DNA scaffold consisting of a DosR binding box (including an upstream central a site and a downstream proximal b site) centered around -43.5 site upstream of the transcription start site, a suboptimal −35 element, a consensus −10 element, and a 13 bp transcription bubble as previously described (**Fig. 1*A***) (31-34). Subsequently, the DosR mutant D54E, which exhibits activation effects comparable to the phosphorylated form, was employed to mimic a phosphorylation-activated DosR and to assemble a functional DosR-TAC. For brevity, the phosphorylated D54E mutant is hereafter referred to as DosR. As shown, all of the subunits of Mtb RNAP and DosR were proportionally incorporated into in the purified complex (***SI Appendix*, Fig. S2*A*, S2*B***). Consistent with this observation, RNAP alone displayed no DNA binding activity, whereas in the presence of DosR, electrophoretic mobility shift assays (EMSA) showed the formation of a noticeable larger ternary complex compared to the binary DosR-DNA complex (***SI Appendix*, Fig. S2*C***). These results indicate that the purified DosR facilitates the formation of an enzymatically active DosR-TAC, potentially achieved by recruiting Mtb RNAP to the promoter DNA.

**Fig. 1.**
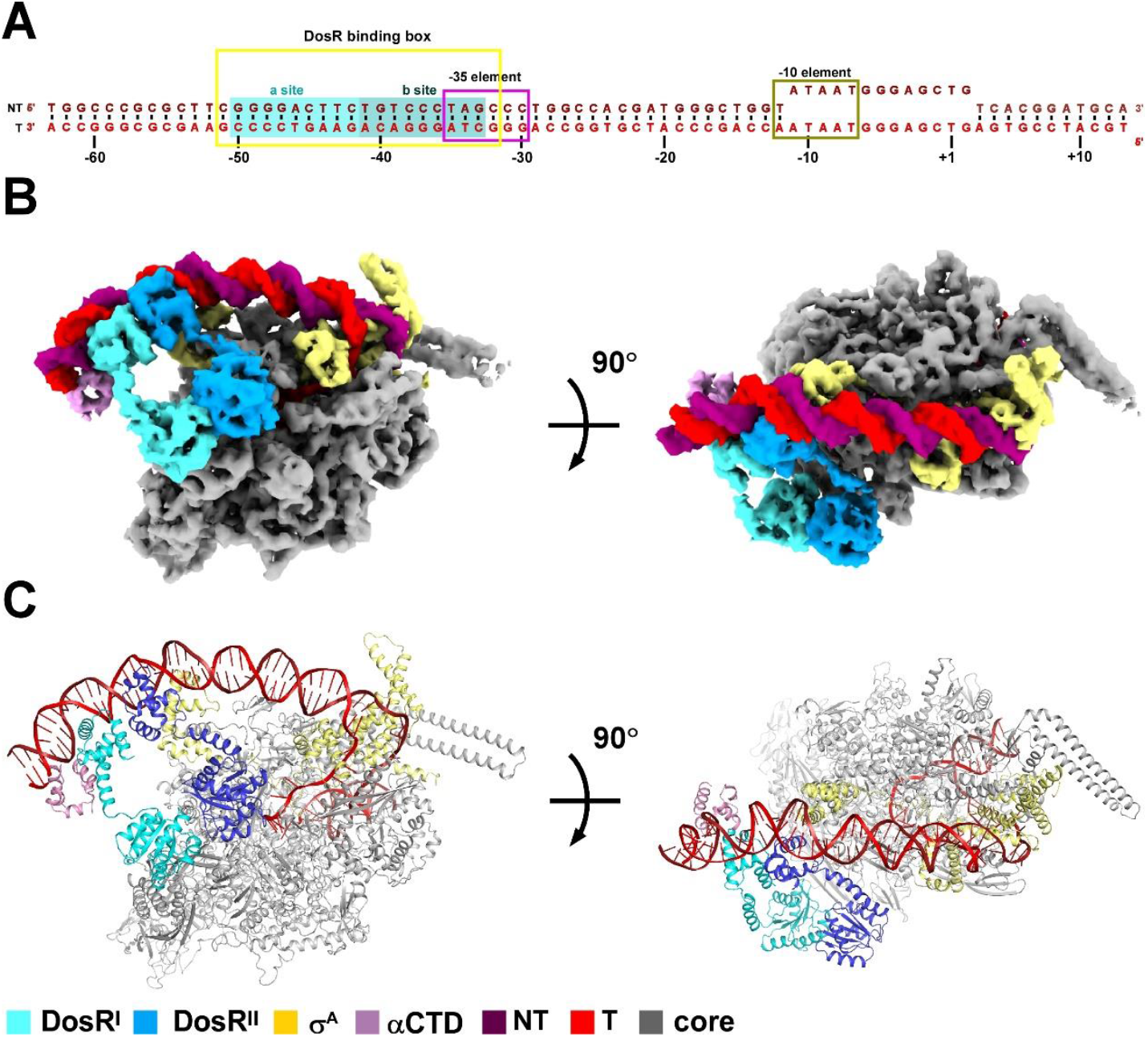
The overall structure of Mtb DosR-TAC. (**A**) DNA scaffold used in structure determination of Mtb DosR-TAC. NT, non-template-strand promoter DNA; T, template-strand promoter DNA. (**B, C**) Two views of the cryo-EM density map (**B**) and structure model (**C**) of Mtb DosR-TAC. The cryo-EM density maps and cartoon representations of Mtb DosR-TAC are colored as indicated in the color key.

Utilizing single-particle reconstruction combined with localized focus refinement targeting the upstream double-stranded DNA and DosR, the cryo-EM map of the DosR-TAC was generated and refined to a nominal resolution of 4.00 Å (**Fig. 1*B*, Table 1, *SI Appendix*, Fig. S3** and **Table S1**). The electron density maps corresponding to the DNA and the multi-subunit Mtb RNAP holoenzyme (α_2_ββ’σ^A^) exhibited high quality, enabling accurate modeling of the downstream DNA scaffold alongside RNAP derived from Mtb RPo (PDB ID: 6VVY) (35). Through successive iterative rounds of local refinement, structural models of the C-terminal domain of Mtb RNAP α subunit (αCTD) and two full-length DosR monomers were effectively fitted into the upstream density, culminating in a comprehensive intact structural model of the Mtb DosR-TAC (**Fig. 1*C***).

**Table 1.**
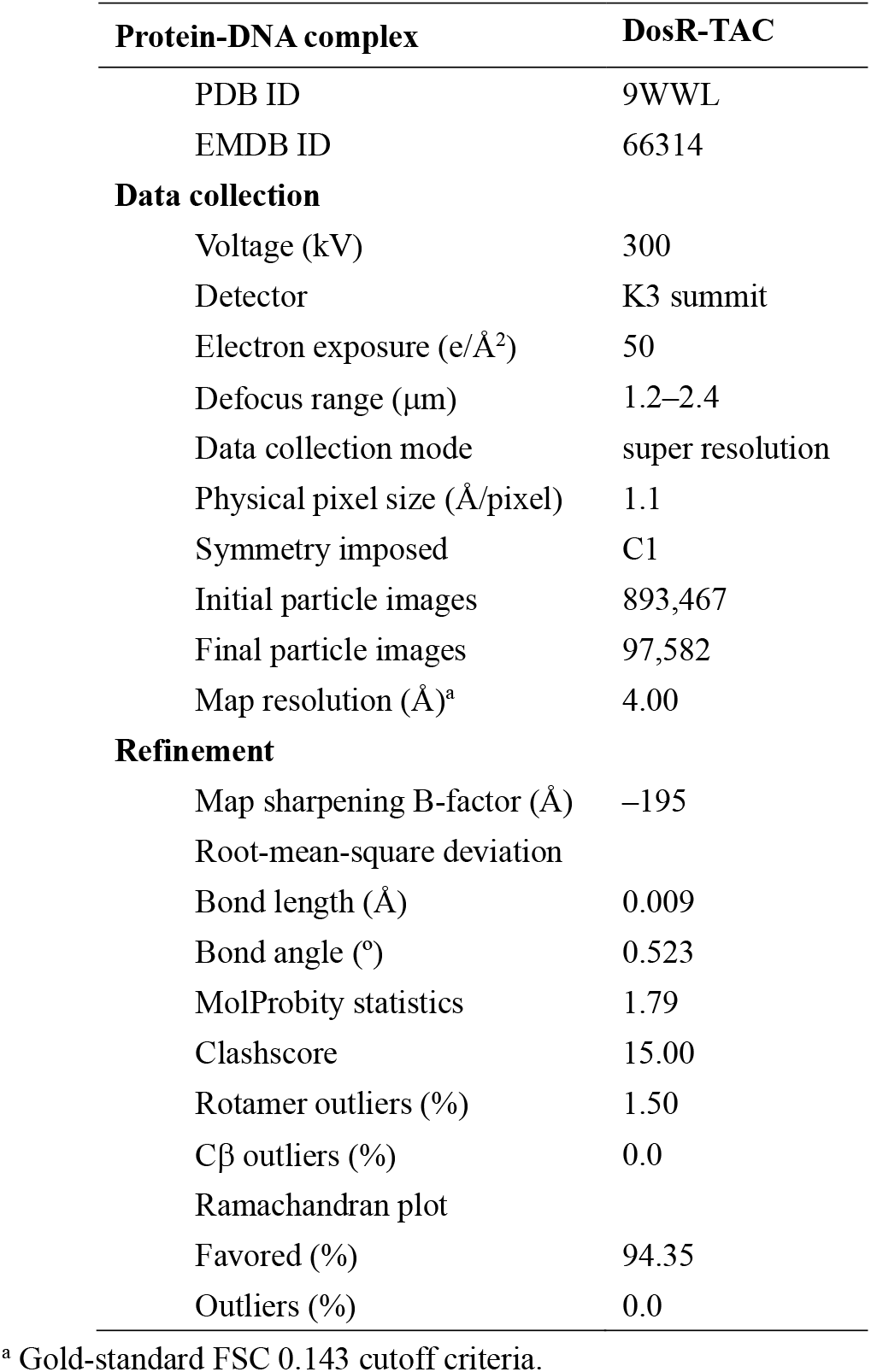
Single particle cryo-EM data collection, processing, and model building for *M. tuberculosis* DosR-TAC.

Notably, within the DosR-TAC, the two DosR monomers (the upstream DosR^I^ and the downstream DosR^II^) assemble into a centrosymmetric dimer that substantially separates its RECs and DBDs via two elongated helical linkers comprising only the α6 helices. This configuration enables the DBDs to induce twisting of the upstream DNA while the RECs engage with the N-terminal domain of Mtb RNAP α subunit (αNTD) (**Figs. 2, 3** and ***SI Appendix*, Fig. S4**). This distinctive structural arrangement displays pronounced conformational deviations from the non-phosphorylated, inactive state of DosR (20), thereby providing novel insights into the conformational flexibility underlying DosR activation and advancing our understanding of the molecular mechanisms governing DosR-dependent transcriptional activation.

**Fig. 2.**
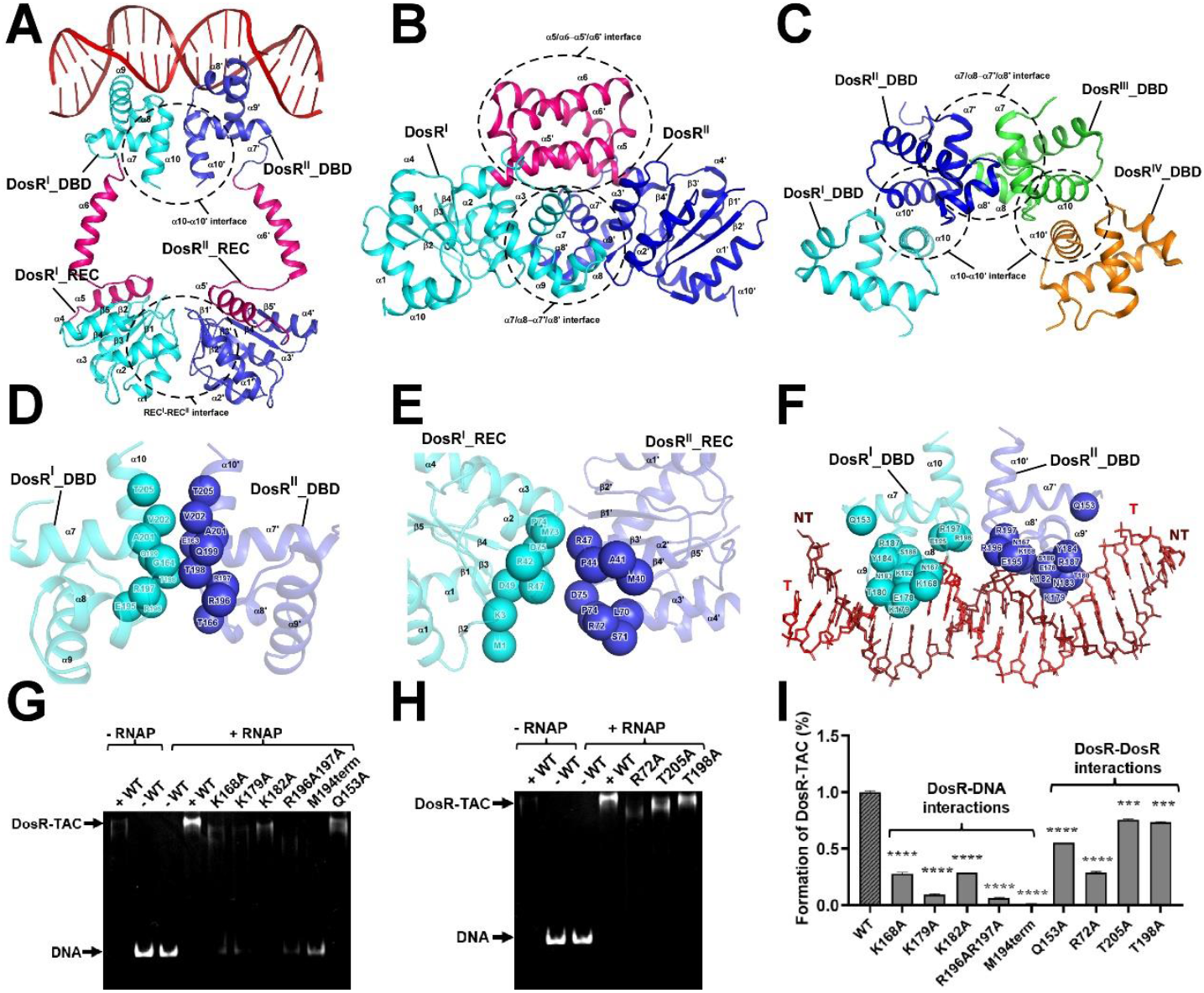
The interactions between two DosR molecules, and the promoter DNA in Mtb DosR-TAC. (A)The relative locations between DosR^I^ and DosR^II^ bound to the promoter DNA in Mtb DosR-TAC. The secondary structural elements involved in DosR^I^ and DosR^II^ are labeled, respectively. (B) Homodimer structure of DosR from PDB entry (3C3W); (**C**) Tetramer structure of DosR_DBD from PDB entry (1ZLJ); (**D, E**) Detailed interactions between Mtb DosR^I^_DBD or DosR^I^_REC and DosR^II^_DBD or DosR^II^_REC. The key residues involved in DosR^I^_DBD or DosR^I^_REC and DosR^II^_DBD or DosR^II^_REC are shown as cyan and blue spheres, respectively. (**F**) Detailed interactions between DosR^I^_DBD and DosR^II^_DBD with promoter DNA. The key residues involved in DosR^I^_DBD and DosR^II^_DBD are shown as green and blue spheres, respectively. (**G-I**) EMSA analyses of DosR-TAC for wild type DosR and the derivatives involved in the DosR-DNA and the DosR-DosR dimeric interfaces. Data for EMSA assays are replicated for 3 times. Error bars represent mean ± SEM of n = 3 experiments.

**Fig. 3.**
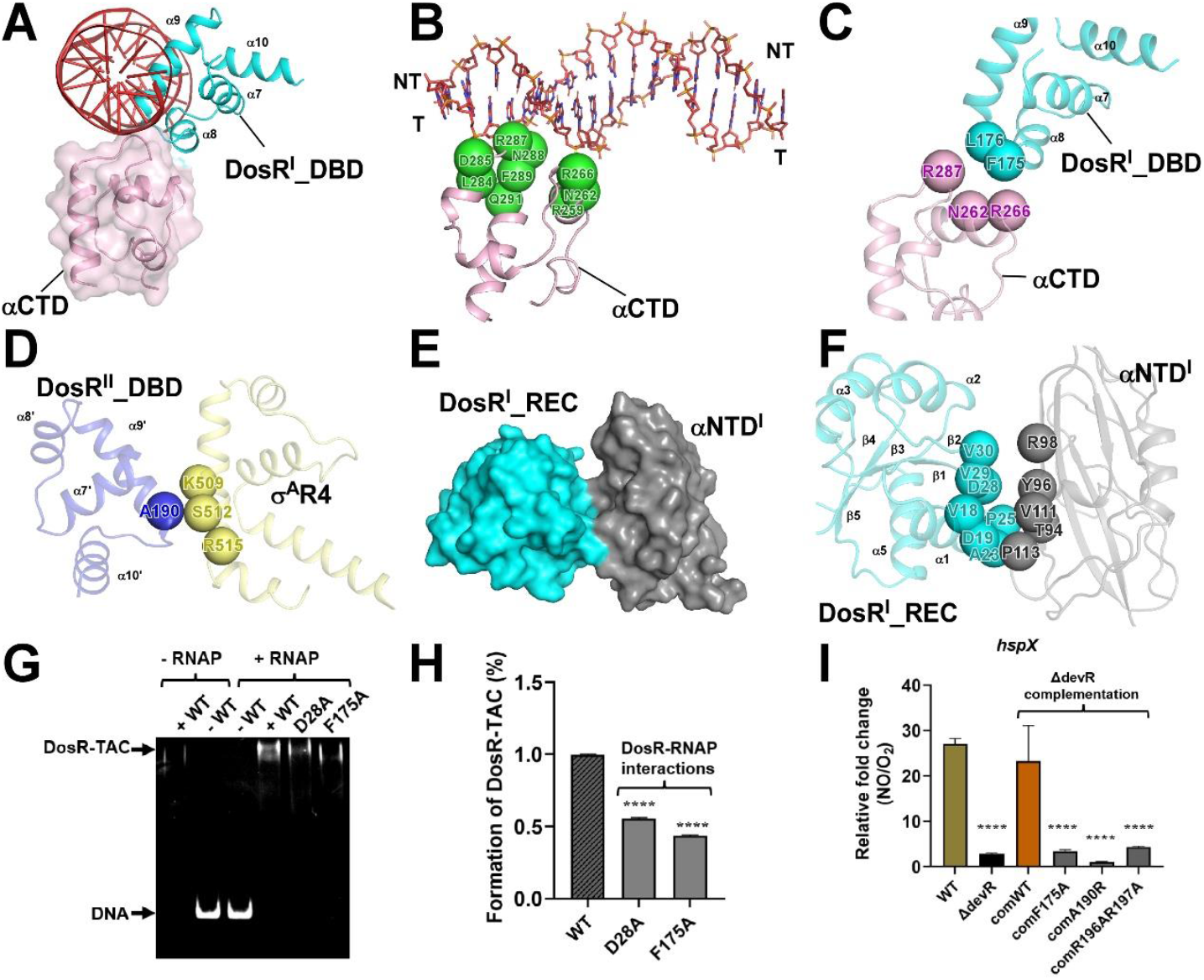
The critical protein-protein interactions between Mtb DosR and RNAP subunit α and σ^A^. (**A**) The relative locations between DosR^I^_DBD and RNAP αCTD bound to the promoter DNA. The RNAP αCTD domain is shown in pink; (**B**) Detailed interactions between DosR^I^ -DBD and RNAP αCTD. The key residues involved are shown as green spheres;(**C**) Detailed interactions between DosR^I^-DBD and RNAP αCTD. Residues involved in interactions between DosR^I^_DBD and RNAP αCTD are shown as cyan and pink spheres, respectively; (**D**) Detailed interactions between DosR^II^ -DBD and RNAP σ^A^R4. Residues involved in interactions between DosR^II^ -DBD and RNAP σ^A^R4 are shown as blue and yellow spheres, respectively; (**E**) Electron density map of the DosR^I^-REC domain and RNAP αNTD domain. The RNAP αNTD domain is shown in gray; (**F**) Detailed interactions between DosR^I^ -REC and RNAP αNTD. The key residues involved in DosR^I^_REC and RNAP αNTD are shown as cyan and gray spheres, respectively; (**G, H**) EMSA analyses of DosR-TAC for wild type DosR, D28A, and F175A; (**I**) qRT-PCR analyses of *hspX* gene regulated by DosR from wild type *M. smegamatis* MC^2^155, *dosR* knockout mutant, wild type and mutated *dosR* complemented strains. Data for EMSA and qRT-PCR assays are means of 3 technical replicates. Error bars represent mean ± SEM of n = 3 experiments.

### Activated DosR undergoes significant conformational changes to afford dimerization and DNA engagement

In contrast to the tetrameric configurations identified in the crystal structures of full-length unphosphorylated DosR, DosR_DBD, and the cocrystal structure of DosR_DBD complexed with promoter DNA (20, 26), two activated DosR monomers undergo significant conformational changes to assemble into an extended centrosymmetric dimer and facilitate specific interactions with the promoter DNA (**Fig. 2** and ***SI Appendix*, Fig. S4**).

In DosR-TAC, the extended DosR dimer is stabilized by both the DosR_RECs and the DosR_DBDs (**Fig. 2*A***). Notably, the original linker helix α5’ in the full-length unphosphorylated DosR structure dissociates from helix α6 and isomerizes into a β5 (comprised residues 98 to 105) and a shorter helix α5 (comprised residues 108 to 120) (***SI Appendix*, Fig. S1** and **Fig. 2, *A-E***). This allosteric effect may enhance the interaction between Q199 and the catalytic D54 residue of the DosR_REC, cause helix α4 to disrupt its interaction with the DosR_DBDs, enhance the interaction between Q199 and the catalytic D54 residue of the DosR_REC, thereby isomerizing DosR_RECs into a canonical (βα)5 fold as observed in other response regulators (20, 33). As depicted in DosR-TAC, the dimeric α10-α10 interface within the DosR_DBDs is mediated by polar and hydrophobic interactions similar to those observed in the AB or CD type dimer of the crystal structure of DosR_DBDs or the cocrystal structure of DosR_DBDs and DNA (**Fig. 2*D***) (20, 26). Besides, a novel dimeric interface between the DosR_RECs is captured for the first time, potentially being maintained by electrostatic interactions (**Fig. 2*E***), presenting a comparable interface gap volume of ∼1300 Å^3^ to the reported DosR_DBD dimers. Consequently, this transformation disrupts the initially interconnected linker helices α5’α6 in the unphosphorylated DosR, causing a separation between the DosR_DBDs and the DosR_RECs. This separation facilitates the dimerization of the DosR_DBDs through the pivotal α10-α10 interface, enabling insertion into the DNA major groove via the helices α9 (**Fig. 2*F***), thereby promoting the formation of an active DosR dimer.

Structural analysis reveals that the activated DosR specifically reads out promoter DNA through the critical residues K179, K182, and N183 located within the conserved helix α9, especially at the GGGACT and TGTCCC motifs, where the DosR dimer induces about ∼40° twist in the upstream DNA. This conformational alteration is facilitated by potential polar interactions and van der Waals forces between the residues (Q153, N167, K168, T180, and Y184 from helices α7-α9) and phosphates of the DNA backbones (**Fig. 2*G***). The structural depiction of the DosR dimer coincides with the mutagenesis studies (27, 37). Correspondingly, mutations in key residues involved in both the dimerization interfaces (Q153, R72, T198, T205) and the DosR-DNA interface (K168, K179, K182, R196, R197) impaired fraction of the DosR-TAC formation, as assessed in the EMSA (**Fig. 2, *G***-***I***). Notably, truncation of helix α10 (M194term mutant) markedly diminished DNA-binding affinity of DosR and nearly abolished its ability to promote DosR-TAC formation (**Fig. 2, *G*** and ***I***), underscoring the essential role of helix α10 and the critical residues in facilitating DosR dimerization and DNA engagement. These findings are in good agreement with the prior biochemical characterizations and support the helix rearrangement hypothesis (16, 20, 21, 28). Collectively, the data illustrate an allosteric activation mechanism of DosR triggered by phosphorylation, highlighting significant conformational flexibility within the crucial β5α5α6 linker.

### DosR recruits Mtb RNAP through interacting with the conserved domains of αCTD, σ^**A**^**R4, and αNTD**

In response to changing environmental signals, the conserved multi-subunit bacterial RNA polymerases (α2ββ’ωσ) consistently depend on a multitude of transcription activators to finely initiate the expression of downstream stress genes, making transcription activation an important checkpoint in regulating gene expression and bacterial survival (35, 37, 38). The RNA polymerase provides versatile surfaces for transcription regulators to attach and fulfill regulatory functions. Among these, the most widely used ones are the conserved domains of αCTD and σ^A^R4 (34, 38). Nevertheless, the usage of αNTD has been rarely reported.

In DosR-TAC, the extended DosR dimer facilitates the recruitment of Mtb RNAP through extensive interactions with the conserved domains of αCTD, σ^A^R4, and αNTD (**Fig. 3**), which further provide structural evidence for the recruitment roles of DosR as verified in the EMSA (***SI Appendix*, Fig. S1*C***). The αCTD engages the promoter UP element (an AT-rich sequence), via the 287 determinant, which comprises residues I284, D285, R287, N288, F289, and Q293. Additionally, the loop (comprised residues F175 and L176) connecting the α8 and α9 helices of the DosR^I^_DBD also interacts with the conserved 265 determinant (comprised residues R259, N262, and R266) of αCTD (**Fig. 3, *A*-*C***). On the same side of the downstream DNA, residues A190 and K191 from the α9 helix of DosR^II^_DBD are likely to contact the conserved helix of σ^A^R4 (**Fig. 3, *D***). These protein-protein interactions collectively stabilize the 18-base pair DosR binding box through the DosR^I^_DBD-DNA-αCTD and DosR^II^_DBD-DNA-σ^A^R4 sandwich-like arrangements. Remarkably, a previously uncharacterized interface between the DosR^I^_REC and αNTD has been identified (**Fig. 3, *E, F***). This unique interface is stabilized by a cluster of hydrophobic interactions, encompassing a buried surface area of approximately 800 Å^2^, thereby establishing a robust platform for DosR to engage promoter DNA.

Accordingly, the EMSA results indicated that mutagenesis of the key residues (D28 and F175) involved in these interfaces suppressed formation of the DosR-TAC (**Fig. 3, *G*** and ***H***). Consistent with this, the *M. smegmatis* Δ*dosR* strains expressing the DosR mutants (F175A, A190R, and R196A/R197A) failed to induce *hspX* under hypoxia conditions, resulting in comparable defects to the Δ*dosR* mutant strain or the R196AR197A mutant (**Fig. 3*I***). Taken together, these findings underscore the significance of the protein-protein interactions in recruiting RNAP, stabilizing DosR-TAC, and activating the transcription of the DosR target genes.

### Physiological insights into the differential regulation of the two sets of *M. smegmatis* DosR

To elucidate the structural basis underlying the differential hypoxic induction between the functionally active DosR_5244 (MSMEG5244) and the functionally inactive DosR_3944 (MSMEG3944) from *M. smegmatis* (21, 22, 24), we conducted sequence alignment analyses of the Mtb DosR, MSMEG5244, and MSMEG3944 homologues (***SI Appendix*, Fig. S1**), and verified through *in vitro* EMSA experiments (**Fig. 4, *A*** and ***B***) and *in vivo* complementation assays in *M. smegamatis* MC^2^155 (**Fig. 4, *C*** and ***D***).

**Fig. 4.**
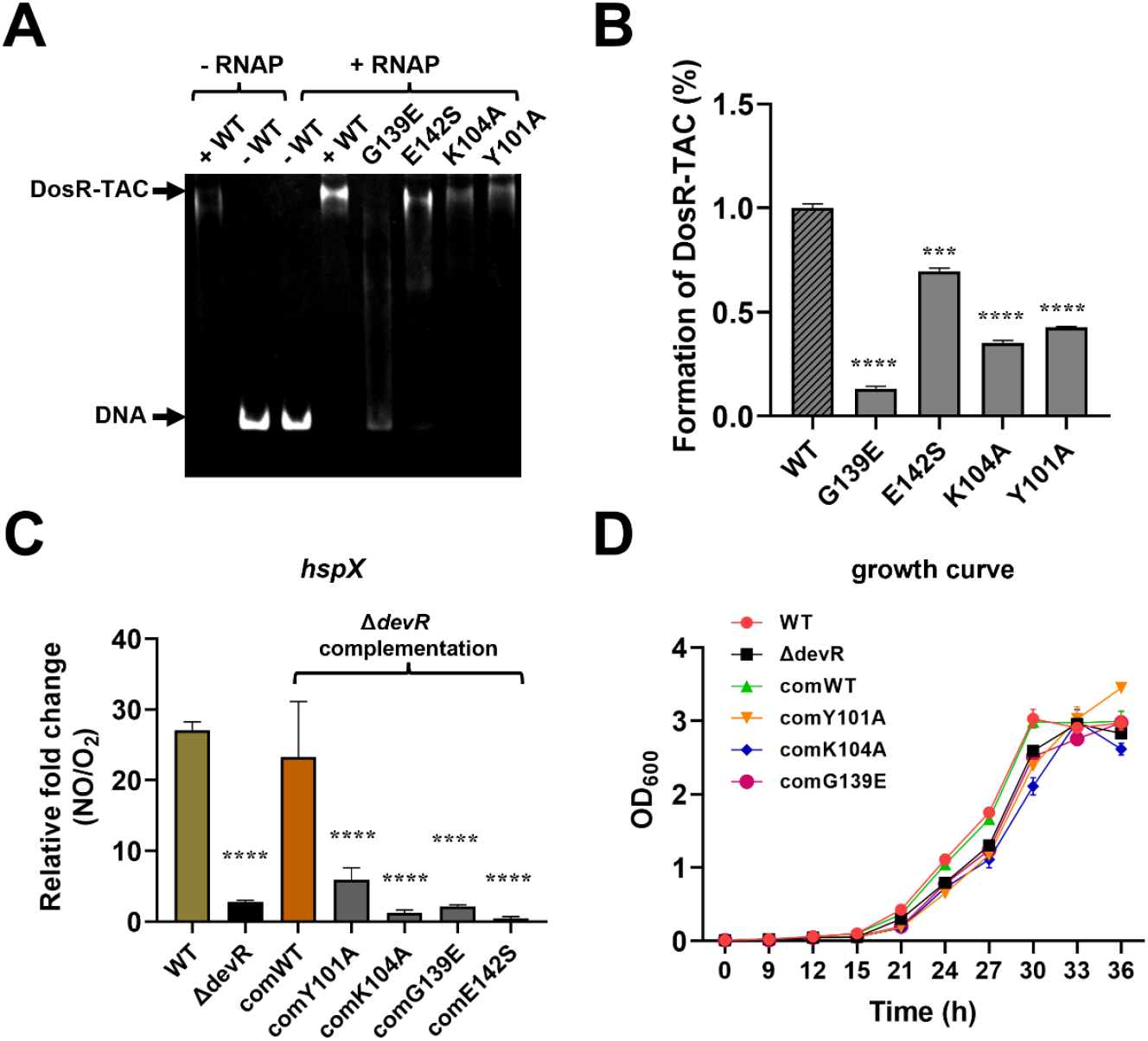
*In vitro* and *in vivo* analyses of DosR regulation. (**A, B**) EMSA analyses of DosR-TAC. (**C**) qRT-PCR analyses of the *hspX* gene regulated by DosR from the wild type *M. smegmatis* MC^2^155, *dosR* knockout mutant, wild type and mutated *dosR* complemented strains. (**D**) Growth curve analysis for wild type *M. smegmatis* MC^2^155 (WT), *dosR* knockout mutant, and the complemented strains of the wild type and mutated *dosR* mutants. Data for the above assays are means of 3 technical replicates. Error bars represent mean ± SEM of n = 3 experiments.

Sequence alignment shows that the functional active DosR_5244 exhibits greater sequence similarity and structural homology with Mtb DosR compared to the functionally inactive DosR_3944, particularly within the conserved DBD domain. However, DosR_3944 displays notable differences in the linker β5α5α6 (***SI Appendix*, Fig. S1**), which may offer novel insights into their divergent functional roles. Consistent with this, site-directed mutagenesis of the differential residues (Y101A, E142S, and G139E) greatly decreased formation of DosR-TAC, with the G139E mutation showing the most pronounced effect (**Fig. 4*A***). Further mutagenesis and complementation assays demonstrated that these mutations nearly abolished the induction of *hspX* under hypoxia conditions (**Fig. 4, *B*** and ***C***). Correspondingly, strains expressing these mutations exhibited slow growth compared to the wild-type (WT) or and the Δ*dosR* strain expressing wild-type *dosR* (**Fig. 4*D***). These results unravel the physiological importance of these residues within the unique linker helices of DosR in mediating hypoxia response. The distinct amino acid composition may induce altered orientations of the linker α6 helix, thereby diminishing the DNA binding affinity of the DosR_DBD in the DosR_3944. This distinctive linker region may thus provide valuable physiological and structural insights for the development of novel therapeutic strategies targeting DosR.

## Discussion

As previously investigated, protein-protein and protein-DNA interactions are essential for transcription activators to hijack bacterial RNAP for initiating specific transcription of stress - responsive genes, enhancing cellular adaptation and pathogenic infections (34, 38). Identifying the key regulatory components that govern these processes is essential for advancing the development of strategies aimed at preventing future pathogenic infections. DosR is among the most extensively characterized response regulators in Mtb, playing a pivotal role in diverse biological processes including dormancy, persistence, and drug tolerance (13, 26). As the essential regulator mediating the adaptation response to hypoxia and nitric oxide stresses of Mtb, considerable efforts have been devoted to elucidating the mechanisms underlying the DosR-dependent transcription activation. In the present study, we successfully resolved the intact cryo-EM structure of the functional DosR-TAC, revealing remarkable conformation flexibility of the key dormancy regulator DosR and providing novel insights into the molecular basis underlying DosR-dependent transcription activation.

By comparing the structure of the unphosphorylated full-length DosR, four distinctive features in DosR-TAC have been identified that contribute to the conformation flexibility and recruitment mechanism of DosR. Firstly, the dimeric linker formed by the α5’ and α6 helices in the unphosphorylated DosR is disrupted by the α5 helix, which promoter the DosR_REC allosterically isomerizes into the canonical (βα)5 fold upon phosphorylation. This rearrangement results in the DosR_RECs and DosR_DBDs being separated solely by the α6 helix. Secondly, the unique α10 helix dissociates from its interaction with the original DosR_RECs and relocates adjacent to the upstream DNA, thereby stabilizing the DosR_DBD-DNA interactions. Thirdly, the two DosR monomers form a centrosymmetric dimer via both the essential α10-α10 interface and the DosR_REC-DosR_REC interface, replacing the original α5’-α6 dimeric and α7-α8 tetrameric interfaces. This extended dimer configuration may establish a distinctive structural architecture for the DosR-type regulators, differentiating them from the canonical transcription activators such as the cyclic AMP receptor protein (CAP) (39), the monomeric global transcription activator SoxS (34), or the tandem OmpR/PhoP family global regulators including PmrA, GlnR, and PhoP (31, 32, 40). Fourth, DosR recruits Mtb RNAP not only through interactions with the conserved αCTD and σ^A^R4 domains, as observed in many transcription activation complexes (34, 38), but also via novel contacts between the DosR^I^_REC and αNTD which provides a stable platform for DosR-TAC. In contrast, the RECs of the nitrate-responsive global transcription factor NarL, which is the DosR closest homologue, are invisible in the cryo-EM structure of NarL-TAC. and there appears no interactions between NarL and the conserved domains of σ^A^R4 and αNTD, as well (33). Although the RECs of the homologous response regulator PmrA are modelled in the cryo-EM structure of PmrA-TAC, the tandem dimeric PmrA interacts with the σ^A^R4, β, β′ subunits instead of αNTD of RNAP (40). As collaboratively suggests diverse cooperative regulation modes of transcription activators to drive RNAP for efficient transcription initiation, and unravel new regulatory roles of RNAP αNTD.

Drawing upon our structural, biochemical, and genetic data, along with prior identifications (13, 17, 20, 26, 41), we propose a comprehensive ‘allosteric activation-recruitment’ model for the DosR-dependent transcription activation mechanism (**Fig. 5**). In the absence of phosphorylation-induced activation signals, DosR remains in inhibition conformation in solution, likely existing in an equilibrium between an inactive dimeric and monomeric state. Within this inactive form, helix α10 interacts with the inactive receiver domain REC’ which lacks helix α5, while dimerization of the linker helices α5 and α6 facilitate the inhibitory configuration (**Fig. 5*A***). Upon phosphorylation of residue D49, steric hindrance disrupts the interaction between helix α10 and the inactive REC’, favoring the formation of the helices α10-α10 dimeric interface. This phosphorylation event induces displacement of helix α5 from helix 6, triggering its isomerization into α5 and β5 structures that integrate into the inactive REC’. Concurrently, this promotes dimerization between two RECs, collectively converting the inactive REC’ into the canonical activated (βα)5 fold REC. This conformational rearrangement significantly extends the linker helix α6, enabling helix α9 from the dimeric DBDs to specifically recognize specific DNA bases (**Fig. 5*B***). Simultaneously, these structural changes facilitate interactions between the upstream and downstream DBDs with αCTD and σ^A^R4, and enable the upstream REC to engage αNTD, supporting recruitment of RNAP and the assembly of an intact stable complex of DosR-TAC (**Fig. 5*C***), thereby enhancing efficient transcription initiation at the downstream DosR target promoters in the presence of nucleoside triphosphates (NTPs) (**Fig. 5*D***). This model provides critical insights into the transcription activation mechanisms of the NarL/UhpA subfamily transcription regulators and potentially other DosR-type response regulators.

**Fig. 5.**
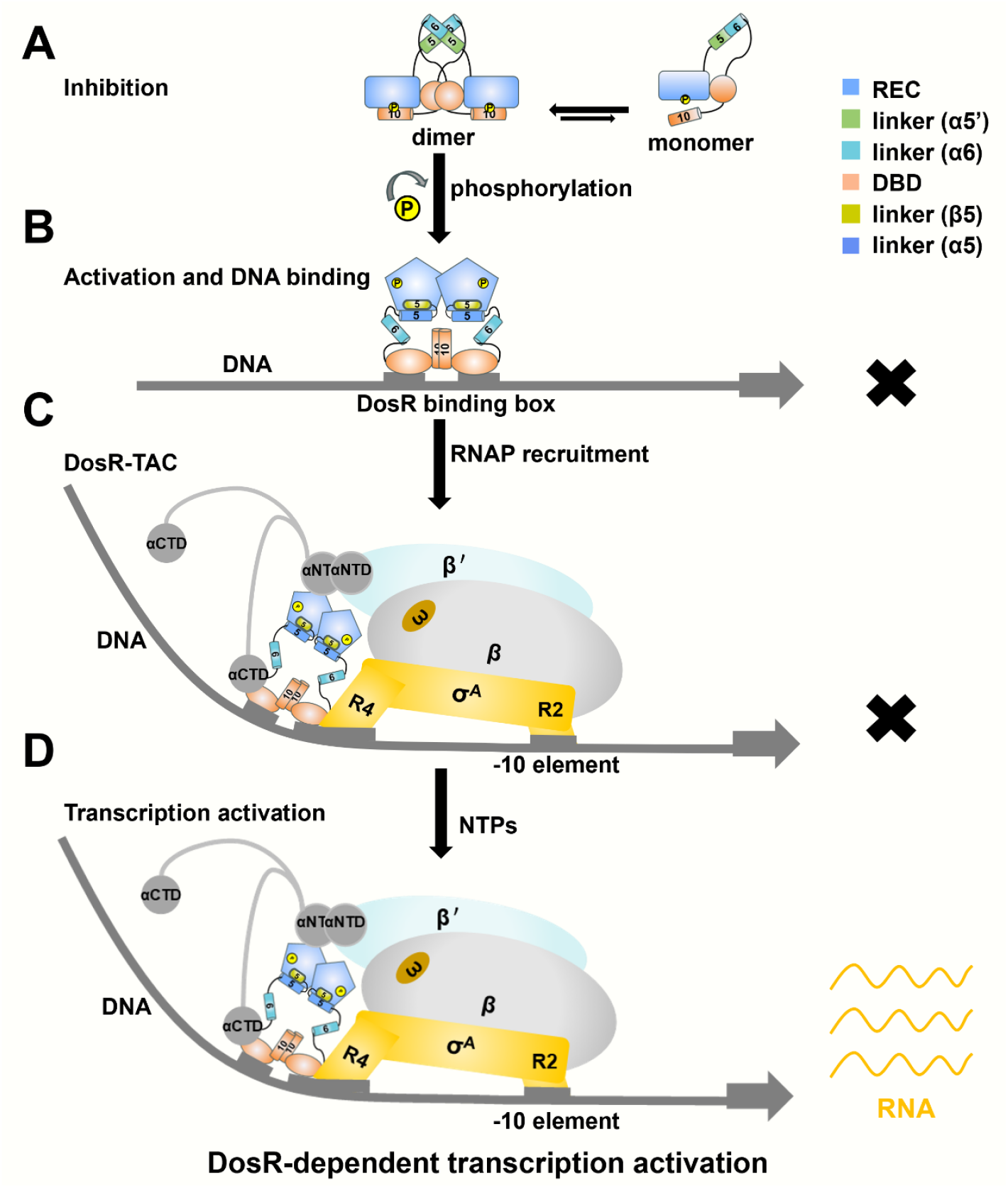
Proposed working model for DosR-dependent transcription activation.

Furthermore, structural comparisons involving the hypoxia responsive Mtb DosR, the *M. smegmatis* DosR_5244, and the functionally irrelevant *M. smegmatis* DosR_3944 reveal the sequence and structural divergences of specific residues that underpin the characteristic conformational flexibility of the critical linker helices. These findings highlight the central regulatory role of this module within DosR homologues and underscore the evolutionary diversity of DosR in transcription activation and stress adaptation. This distinctive linker region may represent a promising target for the development of novel therapeutic agents or interventions aimed at combating dormant Mycobacterium species.

## Materials and Methods

### Plasmids and DNA preparation

To construct the pET28a-*dosR* plasmid, the gene sequence encoding the Mtb DosR was synthesized by Sangon Biotech with a 6-histidine tag at the N-terminal end, and this sequence was subsequently constructed into the pET28a vector containing a T7 promoter. Plasmids with specific amino acid mutations (pET28a-*dosR* derivatives) were constructed using a site-directed mutagenesis kit. To obtain the *hspX* promoter, standard colony PCR method was employed, using the Mtb BCG (Bacillus Calmette-Guérin) genome as the DNA template, and amplification was performed using *hsp-F* and *hsp-R* primers. The PCR products were purified using a Vazyme PCR purification kit. All primers were synthesized by Sangon Biotech and listed in ***SI Appendix* Table S2**.

### Purification of Mtb DosR

The pET28a-DosR plasmid and its derivative plasmids were first transformed into BL21 (DE3) expression strains. Positive monoclonal strains were selected and cultured in 100 mL LB medium containing 50 μg/mL kanamycin at 37 °C, and shaken until the bacterial suspension OD_600_ reached 0.6∼0.8. Subsequently, the culture was transferred at a 1:1000 ratio into 1 L of LB medium containing 50 μg/mL kanamycin. When the OD_600_ of the culture reached approximately 0.6∼0.8, protein expression was induced by supplement of a final concentration of 0.5 mM IPTG under 18 °C for 16 hours. The bacterial precipitates were collected by centrifugation (5500 g, 15 min). Then the cells were resuspended in 25 mL of buffer A (20 mM Tris-HCl, pH 8.0, 0.2 M NaCl, 5% glycerol) and lysed using an ATS AH-10013 cell homogenizer (ATS, Inc). After centrifugation at 4 °C (13000 g) for 30 minutes, the supernatant was loaded onto a 5 ml Ni-NTA agarose (Qiagen, Inc.) column pre-equilibrated with buffer A, and successively washed with 25 mL of buffer A containing 25 mM imidazole, and eluted with 30 mL of buffer A containing 200 mM imidazole. After centrifugation, the eluates were applied onto a 120 mL HiLoad 16/600 Superdex 75 chromatography column (GE Healthcare, Inc.) equilibrated and eluted with buffer B (20 mM Tris-HCl, pH 8.0, 200 mM NaCl, 5 mM MgCl_2_, 1mM DTT). After identification by SDS-PAGE, the eluted fractions containing DosR were combined, concentrated, and stored at -80°C. Finally, DosR achieved a yield of 2mg/L with a purity of approximately 95%. The preparation processes for DosR derivatives were the same as described above.

### Purification of Mtb RNAP

The three plasmids pACYC Duet-*rpoA-rpoD*, pCDF-*rpoZ* and pET Duet-*rpoB-rpoC* were co-transformed into *E. coli* BL21(DE3) expression strain to purify Mtb RNAP as described previously (31, 32). A single colony of the resulting positive transformant was inoculated into 100 mL LB broth containing 35 μg/mL chloramphenicol, 100 μg/mL ampicillin, and 50 μg/mL streptomycin and shaken at 37 ℃C for 16 h. Subsequently, 10 mL of the culture was transferred into 1 L of LB broth containing the same antibiotics for amplification and incubated with shaking at 37℃C. When OD_600_ reached about 0.8, 0.5 mM IPTG was added to induce culture and incubated at 16 ℃C for 18 h. The cells were then centrifuged (5000 g; 4 ℃C 15 min), and the cell pellet was resuspended in 20 ml of lysis buffer A, lysed using a cell homogenizer (ATS, Inc), and centrifuged at 13000g at 4 ℃C for 30 min. The supernatant was precipitated with polyethyleneimine at a ratio of 0.7% (m/v), washed three times with buffer C (10 mM Tris-HCl, pH 7.9, 0.5 M NaCl, 1 mM EDTA, 5% glycerol), extracted with buffer D (10 mM Tris-HCl, pH 7.9, 1 M NaCl, 1 mM EDTA, 5% glycerol), and then precipitated with ammonium sulfate at a ratio of 30.0% (m/v). The precipitate was resuspended in buffer A, and the supernatant was loaded onto a 10 mL Ni-NTA agarose (Qiagen, Inc.) column equilibrated with buffer A, washed with 50 mL buffer A containing 20 mM imidazole, and eluted with 50 mL buffer A containing 200 mM imidazole. The eluent was diluted with buffer E (20mM Tris-HCl, pH 7.9, 5% of glycerol, 1 mM EDTA and 1mM DTT) and loaded onto a Mono Q 10/100 GL column (GE Healthcare, Inc.), and the protein was eluted under a linear concentration gradient of 0.3-0.5 M NaCl. The protein was then equilibrated with buffer F (20 mM Tris-HCl, pH 7.9, 75 mM NaCl, 5 mM MgCl_2_, 1 mM DTT) and loaded onto a 120mL HiLoad 16/600 Superdex 200 column (GE Healthcare, Inc.). The components containing Mtb RNAP were concentrated and stored at -80℃C. The yield was about 0.5 mg/L, and the purity was 95%.

### Assembly of Mtb DosR-TAC

The DNA oligonucleotides used for assembling the Mtb DosR–TAC were synthesized by Sangon Biotech, Inc. The template strand DNA (*hspX* scaffold_T) and nontemplate strand DNA (*hspX* scaffold_NT) of this *hspX* scaffold (**Fig. 1** and ***SI Appendix* Table S2**) were firstly dissolved in nuclease-free water to 100 mM and annealed in a 1:1 ratio in annealing buffer (10 mM Tris-HCl, pH 7.9, 0.2 M NaCl). The Mtb RNAP, *hspX* scaffold and Mtb DosR were then incubated at a molar ratio of 1: 1: 20 at 4 ℃C overnight to assemble the Mtb DosR-TAC. The sample was loaded onto a 120-mL Hiload 16/600 Superdex 200 column equilibrated with buffer F the next day and the column was eluted with the same buffer. The eluent was identified by SDS-PAGE and Electrophoretic mobility shift assays (EMSA). Finally, the fractions containing correctly assembled Mtb DosR-TAC were concentrated to 17 mg/mL using an Amicon Ultra centrifugal filter (10 kDa MWCO, Merck Millipore, Inc.).

### Cryo-EM grid preparation

A final concentration of 4 mM CHAPSO was added to freshly prepared Mtb DosR-TAC samples. After glow-discharge treatment of the Quantifoil grids (R1.2/1.3 Cu 300 mesh holey carbon grids; Quantifoil, Inc.) at 15 mA for 120 s, 3 μL of sample was applied onto the grid and blotted via a Vitrobot Mark IV (FEI) at 10 ℃C and 95% humidity, followed by immediate vitrification in liquid ethane precooled with liquid nitrogen to complete vitrification freezing. The frozen grids were stored in liquid nitrogen until data collection.

### Cryo-EM data collection and processing

Cryo-electron microscopy data collection was conducted on a 300 kV FEI Titan Krios transmission electron microscope equipped with a K3 direct electron detector and a Quantum GIF energy filter. A total of 1956 micrographs were automatically collected through EPU software, at a magnification of 64000x, with a physical pixel size of 1.1 Å, a defocus range of -1.2 to -2.4 μm, and a total electron dose of 50 e^−^/Å^2^. In data processing, MotionCor2 (42) was first used to align and sum subframes, and the contrast transfer function (CTF) parameters of each image were calculated using CTFFIND4 (43). From the summed images, approximately 893, 467 particles were picked with blob picker, template picker and subjected to iterative 2D classification in CryoSPARC v4.2 (44). The automatically picked particles were manually verified, and a second 2D classification was performed to eliminate classes with poor density. By removing the poorly populated classes, 893,467 particles were subjected to hetero refinement by using ab-initio reconstruction and a map of *M. tuberculosis* RPo (PDB ID: 6VVY) low-pass filtered to 40 Å resolution as a reference. After importing class 2 for masked 3D classification on the upstream DosR binding regions, homogeneous refinement, non-uniform refinement, CTF-refinement, local resolution estimation, and local filtering were performed to generate the final density maps by using CryoSPARC v4.2 and UCSF chimera (45). The final 97, 582 particles were further subjected to particle subtraction to keep the signal from the upstream DosR binding regions, accompanied by masked local refinements to enhance the map quality and interpretability. A gold-standard Fourier shell correlation analysis indicated a mean map resolution of 4.0 Å of Mtb DosR-TAC (**Table 1, *SI Appendix* Fig. S3** and **Table S1**).

### Electrophoretic mobility shift assay

Electrophoretic mobility shift assays were performed in buffer (40 mM Tris-HCl, pH 8.0, 100 mM NaCl, 10 mM MgCl_2_, 5% glycerol). The composition and final concentration of the reaction mixture (20μL) were: 16 μM Mtb DosR (or DosR derivative), 100 nM Mtb RNAP, 30 nM *hspX* promoter DNA. The RNAP was preincubated with DNA at 37 ℃C for 10 min, followed by incubation with DosR or DosR derivative at 37 ℃C for 10 min. After adding heparin to a final concentration of 0.03 mg/ml and incubating at 22 ℃C for 2 min, the reaction mixture was subjected to electrophoresis on 5% polyacrylamide slab gel (29:1 acrylamide/bisacrylamide) in 90 mM borate, pH 8.0 and 0.2 mM EDTA, and stained with 4S Red Plus Nucleic Acid Gel Stain (Sangon Biotech, Inc.) following the manufacturer’s procedure. The fraction of DosR-TAC formed by supplement of DosR or its DosR derivatives were repeated for three times and quantified using the Image J software.

### Construction of *M. smegmatis* Δ*dosR* strain

The *M. smegmatis* MC^2^155 (Msm) Δ*dosR* strain was generated via allelic exchange using the pYUB854 plasmid. Briefly, approximately 1500 bp upstream and downstream homology arms of the *dosR* gene were PCR-amplified and gel-purified. These arms were cloned flanking the hygromycin resistance gene (hyg) in the pYUB854 vector. The resulting construct was transformed into *E. coli* DH5α, recovered, and plated on LB agar containing hygromycin (100 μg/mL). Positive clones, identified by colony PCR and confirmed by sequencing, were selected. The verified plasmid was then electroporated into wild-type *M. smegmatis* (Msm WT). Transformants were selected on 7H10 plates containing 50 μg/mL hygromycin, and successful gene deletion was confirmed by colony PCR and sequencing.

### Complementation assay and growth curve assay

For genetic complementation, the *dosR* gene was PCR-amplified and cloned into the integrative plasmid pMV361. The recombinant plasmid (pMV361-*devR*) was transformed into *E. coli* DH5α, extracted, and sequenced. Site-directed mutagenesis was performed using PCR primers designed to introduce the specific point mutation and verified by sequencing. The plasmids were electroporated into the Δ*dosR* strain, selected on 7H10 plates containing 50 μg/mL hygromycin and 20 μg/mL kanamycin, and verified by colony PCR and sequencing. For growth assay, strains were inoculated into 20 mL 7H9 medium (in 150-mL flasks) at an initial OD600 of 0.01 and incubated at 37 °C with shaking at 120 rpm. OD_600_ value was measured every 3 h to obtain growth curves.

### Induction of dormancy

Single colonies were inoculated into 7H9 broth and grown to mid-log phase with OD600 reached to approximately 0.2∼0.3. For induction dormancy program (46), the nitric oxide donor DETA NONOate was added to 10 mL aliquots of culture (in a 15-mL tube) at a final concentration of 100 μM (47). Tubes were tightly capped, sealed with parafilm, and incubated statically at 37 °C for 1 h.

### RNA extraction and qRT-PCR analysis

Total RNA was extracted from cultures as described (48) using TrizoL and treated with DNase. cDNA was synthesized using the High capacity RNA-to-cDNA kit (Applied Biosystems). Quantitative PCR (qPCR) was performed using Power SYBR Green PCR Master Mix (Applied Biosystems) on an applied biosystems quantstudio 5 real-time PCR system. Gene expression levels were normalized to the housekeeping gene *sigA* and expressed as fold change relative to the untreated control.

### Qualification and statistical analysis

To calculate Fourier shell correlations (FSC) of Mtb DosR-TAC, the FSC cut-off criterion was set to 0.143 (49) (***SI Appendix* Fig. S3**). The local resolution of the cryo-EM map was estimated using blocres (50). PHENIX (51) was also used for quantification and statistical analyses during model refinement and validation of Mtb DosR-TAC. Data for EMSA, qRT-PCR, and growth curve assays are replicated for 3 times. Error bars represent mean ± SEM of n = 3 experiments.

## Data Availability

The cryo-EM map and model coordinates have been deposited under the accession numbers: EMDB: 66314 and PDB: 9WWL for *M. tuberculosis* DosR-TAC, EMDB: 66333 for focusing on the upstream DosR binding region in *M. tuberculosis* DosR-TAC. All data are available through the text or in the supplementary file.

## Acknowledgements

We appreciate Liangliang Kong, Guangyi Li, Yun Song, and Jialin Duan of the Electron Microscopy System at the National Facility for Protein Science in Shanghai (NFPS), Shanghai Advanced Research Institute, Chinese Academy of Sciences, and Shenghai Chang at the Center of Cryo-Electron Microscopy in Zhejiang University School of Medicine for providing technical support and assistance in data collection. We thank the Experiment Center for Science and Technology, Nanjing University of Chinese Medicine for experimental assistance. We thank the Core Facilities, Zhejiang University School of Medicine for technical support.

This work was funded by the National Natural Science Foundation of China (32570222, 32270037, 32270192, 82311530689), the Jiangsu Qinglan Project to J.S., the Natural Science Foundation of Jiangsu Province (SBK2023030145, BK20211302), the National Key R&D Program of China (2023YFC2308200), the landmark talent training project of Nanjing University of Chinese Medicine (RC202404), the special fund project of Nanjing Drum Tower hospital for the transformation of scientific and technological achievements (202404), the 2025 Graduate Student Scientific Research and Practice Innovation Program of Nanjing University of Chinese Medicine (035025021001G-103), and the 2025 National Undergraduate Innovation Training Program of the Higher Education Department of the Ministry of Education (202510315023).

## Author Contributions

J. S., ZZ. F., H. F, YY. H., LQ. X., Q. S., and W. C. prepared the proteins, performed the biochemical assays and physiological assays. J. S., ZZ. F., and LQ., X. assembled the cryo-EM samples and collected the cryo-EM data. W. L. and Y. F. performed the cryo-EM structure determination. J. S., W. L., Y. F., and LD. L. designed the study. All authors contributed to data analysis. J. S., ZZ. F. and W. L. wrote the paper with the help from all the coauthors.

