## Supplemental figures and tables for "Molecular mechanism of the key dormancy regulator DosR-dependent transcription activation in *Mycobacterium tuberculosis*"

**This PDF includes:**

**Figures S1 to S5**
**Tables S1 to S2**
**SI References**

### Supplemental Figures

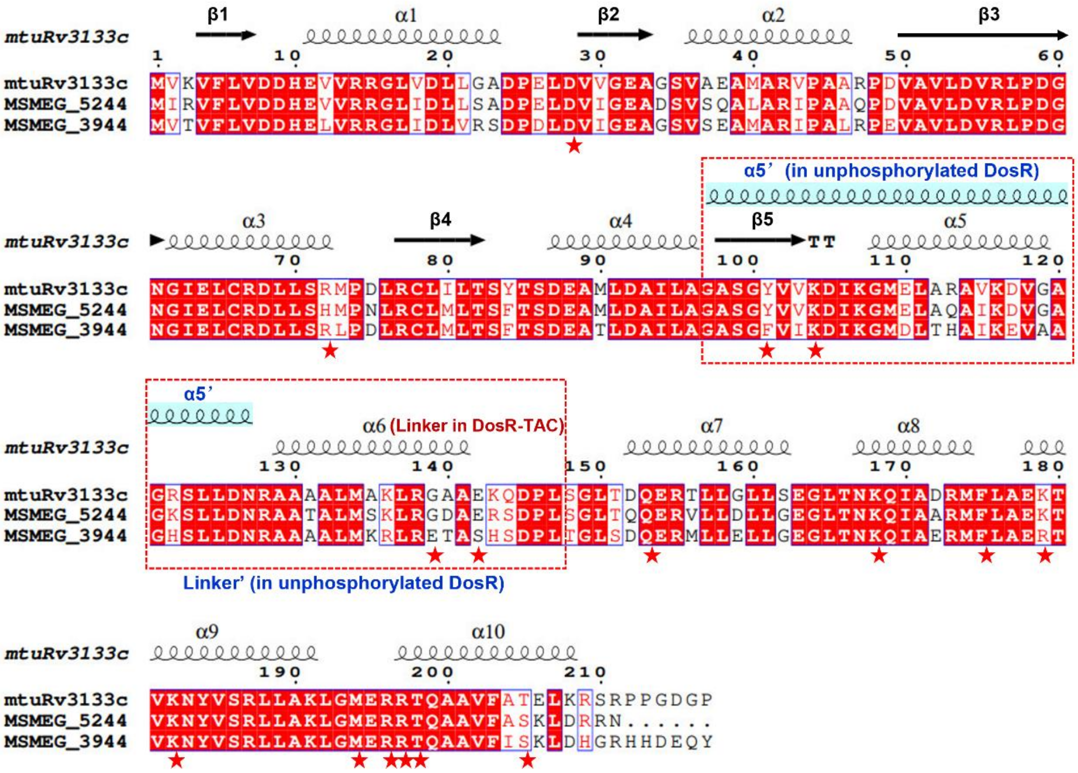


**Fig. S1 Structure based sequence alignment of *M. tuberculosis* DosR and its orthologues in *M. smegamatis*.**

Sequences alignment of *M. tuberculosis* (Mtb) DosR (mtuRv3133c), *M. smegamatis* DosR_5244 (MSMEG_5244), and DosR_3944 (MSMEG_3944). The secondary structure elements of Mtb DosR are shown at the top and as the template in the ESpript Web server (1). The invariant residues are highlighted in red, and the conserved residues are boxed. The helix α6 Linker in DosR-TAC, helix α5' and Linker' in the unphosphorylated DosR structure (PDB ID: 3C3W) are indicated and shown with shaded cyan color, boxed with red dotted line, respectively. The residues mutated in the text regarding the DosR-DNA, DosR-RNAP interfaces and the differentiated linker (spanning from β5 to helix α6) are indicated by red stars, respectively.

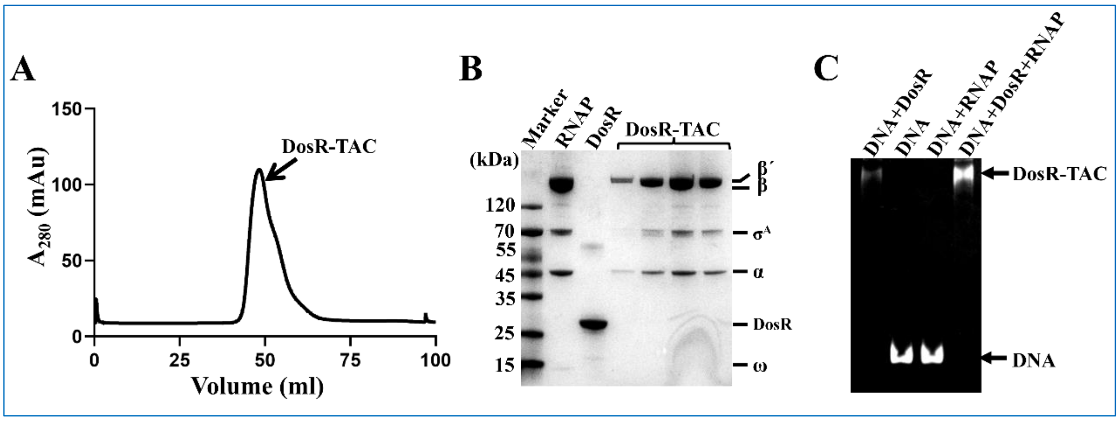


**Fig. S2 Purification and verification of Mtb DosR-TAC.**

**(A)** Gel purification map of Mtb DosR -TAC analyzed by a 120mL HiLoad 16/600 Superdex 200 column. **(B)** SDS-PAGE of the purified Mtb DosR-TAC. **(C)** Formation of Mtb DosR-TAC on the *hspX* promoter as determined by the EMSA experiment. Reaction condition is described in the *Materials and Methods*. Bands of free DNA and DosR-TAC are indicated by arrows on the right, respectively.

56

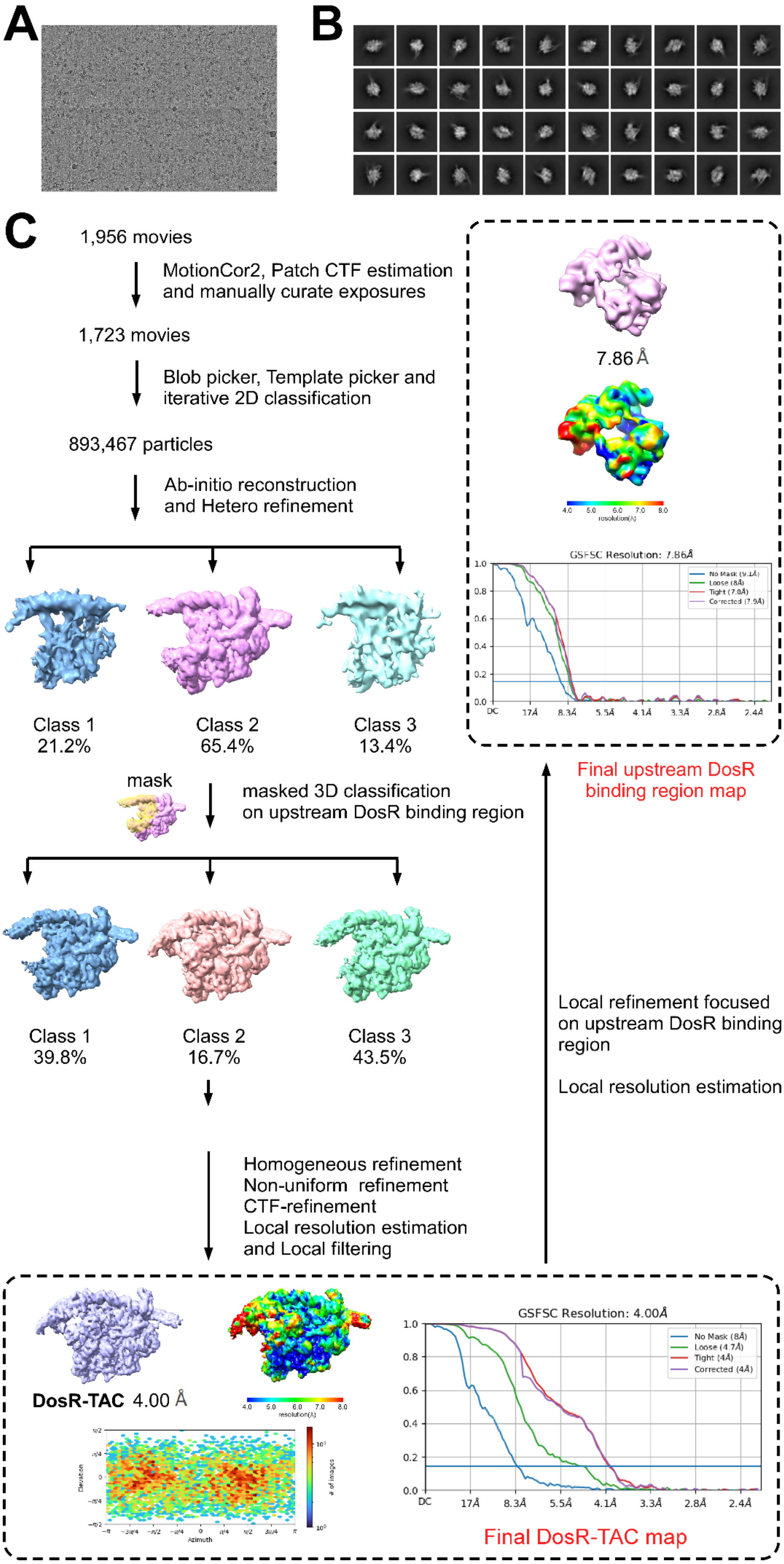

A
B
C
1,956 movies
MotionCor2, Patch CTF estimation
and manually curate exposures
1,723 movies
Blob picker, Template picker and
iterative 2D classification
893,467 particles
Ab-initio reconstruction
and Hetero refinement
Class 1
21.2%
Class 2
65.4%
Class 3
13.4%
mask
masked 3D classification
on upstream DosR binding region
Class 1
39.8%
Class 2
16.7%
Class 3
43.5%
Homogeneous refinement
Non-uniform refinement
CTF-refinement
Local resolution estimation
and Local filtering
7.86 Å
GSFSC Resolution: 7.86Å
No Mask (9.1Å)
Loose (8Å)
Tight (7.8Å)
Corrected (7.9Å)
Final upstream DosR
binding region map
Local refinement focused
on upstream DosR binding
region
Local resolution estimation
DosR-TAC 4.00 Å
GSFSC Resolution: 4.00Å
No Mask (8Å)
Loose (4.7Å)
Tight (4Å)
Corrected (4Å)
DC
17Å
8.3Å
5.5Å
4.1Å
3.3Å
2.8Å
2.4Å
Azimuth
Elevation
Final DosR-TAC map

**Fig. S3 Cryo-EM image processing flow chart of Mtb DosR-TAC.**

**(A)** Representative raw cryo-EM image of Mtb DosR-TAC. **(B)** 2D classes of Mtb DosR-TAC. **(C)** Data processing pipeline for Mtb DosR-TAC. Combination of the 3D refinements on RNAP main body and focusing on the upstream DosR region by using all the particles picked from good 3D classes generated a 4.00 Å map of Mtb DosR-TAC as analyzed in CryoSPARC (2). Angular distribution plot, the final maps, half-map FSC curves (3), and accompanying local resolution illustrations are enclosed in the dashed black boxes.

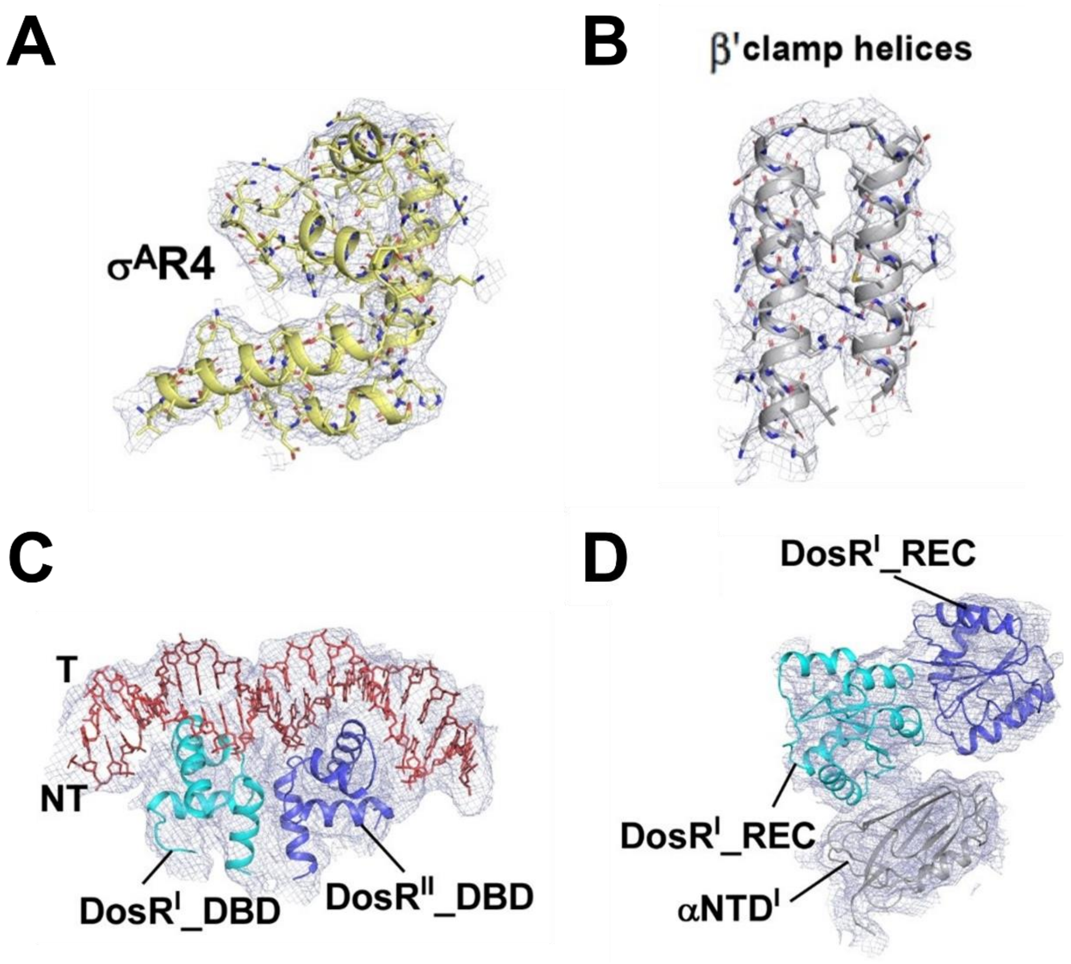


**Fig. S4 Representative cryo-EM densities of superimposed models in Mtb DosR-TAC.**

(**A**) Cryo-EM density map (blue mesh) and the superimposed model of $\sigma^{A}$R4; (**B**) Cryo-EM density map (blue mesh) and the superimposed model of β' clamp helices; (**C**) Cryo-EM density map (blue mesh) and the superimposed model of $\text{DosR}^{I}$_DBD, $\text{DosR}^{II}$_DBD and the promoter DNA; (**D**) Cryo-EM density map (blue mesh) and the superimposed model of $\text{DosR}^{I}$_REC, $\text{DosR}^{II}$_REC and $\alpha\text{NTD}^{I}$. Other colors are shown as in Fig. 1.

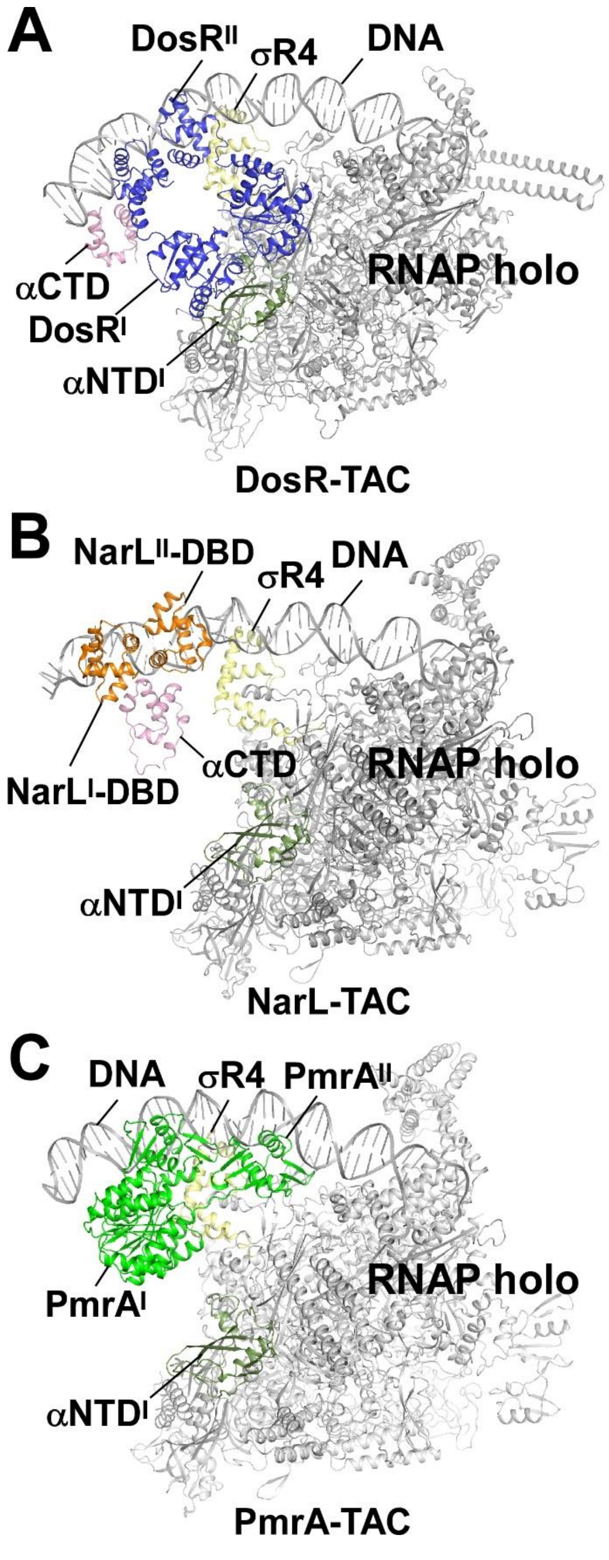


**Fig. S5** Structural comparisons of Mtb DosR-TAC, NarL-TAC (PDB ID:8U3B) (4), and PmrA-TAC (PDB ID:8JO2) (5). Blue, DosR; orange, NarL; green, PmrA; dark gray, $\sigma^A$ RNAP αNTD, αCTD, β and β'.

### Supplemental Tables

**Table S1. Single particle cryo-EM data collection, processing for focusing on DosR region in *M. tuberculosis* DosR-TAC.**

| **Protein-DNA complex** | **DosR region in DosR-TAC** |
|---|---|
| EMDB ID | EMD-66333 |
| **Data collection** | |
| Voltage (kV) | 300 |
| Detector | K3 summit |
| Electron exposure (e/Å$^2$) | 50 |
| Defocus range (μm) | 1.2–2.4 |
| Data collection mode | super resolution |
| Physical pixel size (Å/pixel) | 1.1 |
| Symmetry imposed | C1 |
| Initial particle images | 893,467 |
| Final particle images | 22,425 |
| Map resolution (Å)[a] | 7.86 |

[a] Gold-standard FSC 0.143 cutoff criteria.

**Table S2. Primer sequences used in this study.**

| **Primer name** | **Sequence (5′to 3′)** |
|---|---|
| *hspX -F* | GGCGGTTGGCAGACAAC |
| *hspX -R* | ATCCTTCCTTCGTACGTGCCTGCCTCCTAATCGATGGAAAC |
| *hspX* scaffold_T | TGCATCCGTGAGTCGAGGGTAATAACCAGCCCATCGTGGCCAGGGCTAGGGACAGAAGTCCCCGAAGCGCGGGCCATTTGTCCGCGCCCGTCGGTGATCCACTTGGGGACCATTGACCCTGTTG |
| *hspX* scaffold_NT | CAACAGGGTCAATGGTCCCCAAGTGGATCACCGACGGGCGCGGACAAATGGCCCGCGCTTCGGGGACTTCTGTCCCTAGCCCTGGCCACGATGGGCTGGTATAATGGGAGCTGTCACGGATGCA |
| *D28A-F* | CGGAACTGGCTGTTGTTGGTGAAGCG |
| *D28A-R* | CAACAACAGCCAGTTCCGGATCAGCAC |
| *D54E_F* | GGTTCTGGAGGTTCGTCTGCCGGATG |
| *D54E_R* | CAGACGAACCTCCAGAACCGCAAC |
| *R72A_F* | GATCTGCTGAGCGCTATGCCGGATCTGCG |
| *R72A_R* | AGATCCGGCATAGCGCTCAGCAGATCACG |
| *Y101A-F* | GCGCGAGCGGCGCCGTTGTTAAAGATAT |
| *Y101A-R* | TTTAACAACGGCGCCGCTCGCGCCCGC |
| *K104A-F* | GCTACGTTGTTGCAGATATTAAAGGTATGG |
| *K104A-R* | TTTAATATCTGCAACAACGTAGCCGCTC |
| *G139E-F* | GAAACTGCGTGAGGCGGCGGAAAAACAG |
| *G139E-R* | TTCCGCCGCCTCACGCAGTTTCGCCATC |
| *E142S-F* | CGTGGTGCGGCGTCAAAACAGGACCCGC |
| *E142S-R* | GGTCCTGTTTTGACGCCGCACCACGCAG |
| *Q153A_F* | CCTGACCGATGCGGAACGTACCCTGC |
| *Q153A_R* | TACGTTCCGCATCGGTCAGGCCGCTC |

| | |
|---|---|
| *K168A_F* | CTGACCAACGCACAGATCGCGGATCG |
| *K168A_R* | GCGATCTGTGCGTTGGTCAGGCCTTCG |
| *F175A-F* | GATCGTATGGCCCTGGCGGAAAAAACC |
| *F175A-R* | CCGCCAGGGCCATACGATCCGCGATC |
| *K179A_F* | CCTGGCGGAAGCAACCGTTAAAAACTACG |
| *K179A_R* | TAACGGTTGCTTCCGCCAGGAACATACG |
| *K182A_F* | GAAAAAACCGTTGCAAACTACGTTAGCCGTC |
| *K182A_R* | CTAACGTAGTTTGCAACGGTTTTTTCCGCCAG |
| *A190R-F* | GTCTGCTGCGGAAACTGGGCATGGAAC |
| *A190R-R* | CCCAGTTTCCGCAGCAGACGGCTAACG |
| *M194term_F* | GAAACTGGGCTAGGAACGTCGTACCCAG |
| *M194term_R* | ACGACGTTCCTAGCCCAGTTTCGCCAG |
| *R196AR197A_F* | GGCATGGAAGCTGCTACCCAGGCGGCGG |
| *R196AR197A_R* | CTGGGTAGCAGCTTCCATGCCCAGTTTCG |
| *T198A_F* | CATGGAACGTCGTGCCCAGGCGGCG |
| *T198A_R* | CCGCCTGGGCACGACGTTCCATGCC |
| *T205A_F* | GGTTTTCGCGGCCGAACTGAAACGTAG |
| *T205A_R* | TTTCAGTTCGGCCGCGAAAACCGCCGC |
